# Specialization of chromatin-bound nuclear pore complexes promotes yeast aging

**DOI:** 10.1101/2021.06.28.450139

**Authors:** Anne C. Meinema, Theo Aspert, Sung Sik Lee, Gilles Charvin, Yves Barral

## Abstract

The nuclear pore complex (NPC) mediates nearly all exchanges between nucleus and cytoplasm, and changes composition in many species as the organism ages. However, how these changes arise and whether they contribute themselves to aging is poorly understood. We show that in replicatively aging yeast cells attachment of DNA circles to NPCs drives the displacement of the NPCs’ nuclear basket and cytoplasmic complexes. Remodeling of the NPC resulted from the regulation of basket components by SAGA, rather than from damages. These changes affected NPC interaction with mRNA export factors, without affecting the residence of import factors or engaging the NPC quality control machinery. Mutations preventing NPC remodeling extended the replicative lifespan of the cells. Thus, our data indicate that DNA circles accumulating in the mother cell drive aging at least in part by triggering NPC specialization. We suggest that antagonistic pleiotropic effects of NPC specialization are key drivers of aging.

## Introduction

Nuclear pore complexes (NPCs), which mediate transport of cargos between the nucleus and the cytoplasm, undergo substantial changes during aging from yeast to mammals (Rempel et al., 2020). In post-mitotic cells such as neurons, core NPC components are very stable, tend to become oxidized over time and progressively lose functionality (Savas et al., 2012; Toyama et al., 2013). Accordingly, in the neurons of old rats NPCs become increasingly leaky with age, affecting the proper retention of nucleoplasmic proteins in the nucleus (D’Angelo et al., 2009). On the opposite, yeast NPCs are targeted by a quality control machinery that removes damaged or misassembled NPCs (Webster et al., 2014). Accordingly, no oxidized NPCs are observed in replicative aging yeast cells (Rempel et al., 2019). Still, proteomics analysis of the old yeast cells indicate that they progressively change composition, losing their nuclear basket and cytoplasmic complexes (Janssens et al., 2015; Rempel et al., 2019). Similar changes are observed in the liver of aging rats (Ori et al., 2015; Rempel et al., 2020). However, little is known about the underlying mechanisms and whether this process is simply a consequence or contributes to aging. Particularly, the NPC changes observed in replicative aging yeast cells did not lead to leakage but rather to a more stringent compartmentalization of the tested cargos (Morlot et al., 2019; Rempel et al., 2019). Given the pivotal role of NPCs in a wide variety of cellular processes beyond nucleocytoplasmic exchange, from the regulation of gene expression to DNA repair, it is plausible that defects in NPC function could affect cellular physiology and contribute to the ageing process. However, a clear view of how age affects NPCs and their functionality has been lacking so far, precluding a robust understanding of whether and how NPCs indeed contribute to the aging process.

The budding yeast *Saccharomyces cerevisiae* undergoes both post mitotic and replicative aging (Longo et al., 2012). In the first case, also called chronological aging, starved yeast cells progressively lose viability, with their mortality increasing exponentially with time (Fabrizio and Longo, 2003). In the second case, cells that proliferate in rich nutrient conditions divide asymmetrically through the budding of rejuvenated daughters off the surface of their aging mother cell, which retain and accumulate diverse aging factors (Mortimer and Johnston, 1959; reviewed in Denoth Lippuner et al., 2014). After undergoing 20-30 divisions and generating as many daughter cells, these mother cells stop dividing and ultimately die. Although a number of the involved aging factors and some of the mechanisms of their retention in the mother cell have been characterized, little is known about how they affect the viability of old mother cells.

Extrachromosomal DNA circles (ERCs, (Denoth Lippuner et al., 2014; Morlot et al., 2019; Sinclair and Guarente, 1997)), which are generated in virtually all mother cells at some point in their lifespan through excision of one or more rDNA repeat(s) from the rDNA locus, are a particularly prominent aging factor. These episomes lack a centromere and are very efficiently retained in the mother cell at mitosis. As they replicate once per division cycle, they accumulate exponentially in the nucleus of the old mother cell over time. At the end of their life, the mother cell may contain up to thousand ERCs, which represent about as much DNA as the rest of the genome (Denoth-Lippuner et al., 2014; Morlot et al., 2019). Several lines of evidence establish that ERCs accumulation promote aging of the cell. Mutants that form ERCs at a reduced rate, such as the *fob1Δ* mutant cells, show an extended replicative lifespan (RLS; (Defossez et al., 1999). In reverse, cells in which recombination in the rDNA is derepressed, such as cells lacking the sirtuin Sir2, show a higher rate of ERC formation, accumulate them faster and are short-lived (Kaeberlein et al., 1999). Finally, artificially introducing an ERC (Sinclair and Guarente, 1997) or any other replicating DNA circle in a young cell (Denoth- Lippuner et al., 2014) causes premature aging. However, how ERCs promote aging is not known.

The tight retention of the ERCs in the mother cell depends on their attachment to NPCs at the nuclear periphery (Denoth-Lippuner et al., 2014). Attachment is mediated by the SAGA complex, a large multi-subunit complex harboring acetyl-transferase activity (provided by Gcn5) and which binds both NPCs, via the SAGA subunit Sgf73 and the stable nucleoporin Nup1 (Jani et al., 2014; Köhler et al., 2008) and chromatin (Durand et al., 2014; Wang et al., 2020). Since yeast cells undergo a closed mitosis and NPCs remain intact through mitosis, attachment of the DNA circles to NPCs leads to the retention of both in the mother cell. Retention is facilitated by the presence of a ceramide-based, lateral diffusion barrier in the outer-membrane of the nuclear envelope (Baldi et al., 2017; Clay et al., 2014; Denoth-Lippuner et al., 2014; Megyeri et al., 2019; Prasad et al., 2020; Shcheprova et al., 2008). Accordingly, cells that fail to attach ERCs to NPCs (such as the cells lacking the SAGA subunits Sgf73 and Gcn5) or fail to assemble the diffusion barrier (in cells lacking the barrier protein Bud6) cannot tightly confine the ERCs to the mother cell anymore and do not accumulate them over time. Providing further evidence for ERCs promoting cellular aging, these cells and the mutant strains unable to form a lateral diffusion barrier in the nuclear envelope (such as the cells lacking the ceramide synthase Lag1, the sphinganine C4-hydroxylase Sur2) show a dramatically increased longevity (Clay et al., 2014; D’Mello et al., 1994; Denoth Lippuner et al., 2014; Megyeri et al., 2019; Shcheprova et al., 2008). Interestingly, because ERCs and NPCs are retained together in the mother cell during cell division, aging wild type cells also accumulate NPCs and this accumulation is relaxed in cells lacking either of the Fob1, Sgf73 or Bud6 proteins (Denoth-Lippuner et al., 2014). Thus, these accumulating NPCs may contribute to the mechanisms by which ERCs promote aging of the cell.

The nuclear pore complex is a ∼100 megadalton entity comprising repetitions of more than 30 different subunits, called nucleoporins (short Nups)(Beck and Hurt, 2017; Kim et al., 2018). The core of the NPC is inserted in the nuclear envelope and stabilized by protein rings at its periphery. The center is filled by so-called FG-Nups, which both form a barrier to passive diffusion through the pore and facilitate the passage of transport intermediates across the NPC. On their cytoplasmic face, NPCs are decorated by a complex of cytoplasmic Nups which have been also described as fibrils or filaments (Fernandez-Martinez et al., 2016; Strambio-De-Castillia et al., 2010)(Fig. 1A). On their nucleoplasmic side, they assemble the so-called nuclear basket formed mainly of the extended protein TPR, known as Mlp1 and Mlp2 in yeast (Strambio-de-Castillia et al., 1999). Diverse studies have involved the basket in controlling the interaction of chromatin with the NPC (Niepel et al., 2013). However, whether and how DNA circle attachment to NPCs affects these NPCs is not known. Therefore, we have investigated here whether the anchorage of DNA circles to NPCs affects their composition during replicative aging and whether this has an effect on cellular viability.

**Figure 1.**
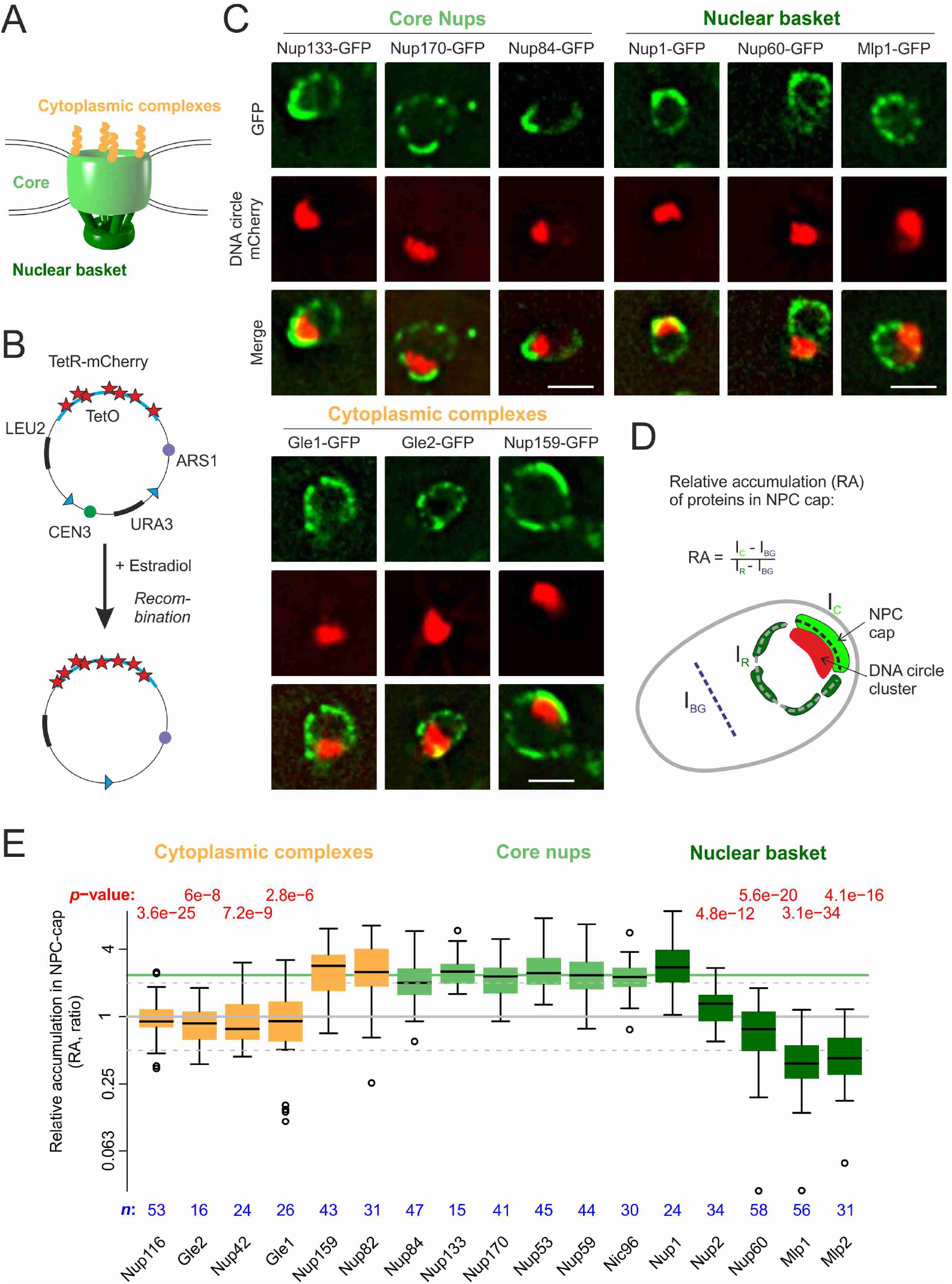
DNA circle anchored NPCs lack the peripheral subunits. (A) Cartoon of the NPC, showing the scaffold core, cytoplasmic complexes and nuclear basket. (B) Cartoon of an engineered DNA circle with excisable centromere (CEN3) and TetR-mCherry decorated TetO repeats. The circle contains an autonomic replication sequence (ARS1) and selection markers (LEU2, URA3). (C) Fluorescent images of nuclei in yeast cells with accumulated DNA circles; the DNA circle clusters are labeled with TetR-NLS-mCherry (red) and different nucleoporins labeled with GFP (green). Scale bar is 2 µm. More images in Figure 1 – figure supplement 1. (D) Cartoon exemplifying the quantification of protein accumulation around the DNA cluster. The NPC cap was selected by a line scan through the nuclear envelope adjacent to the DNA circle cluster. The relative accumulation (RA) is defined as the ratio of background corrected GFP intensity in the NPC cap (I_c_ – I_BG_) over that in the rest of the nucleus (I_R_ – I_BG_). (E) Quantification of GFP-labeled nucleoporin accumulation in the NPC cap localized at the DNA circle cluster. The relative fluorescence intensity in the NPC cap is plotted on a log2-scale. The median accumulation of the core nucleoporins is indicated (green line). *p-*value stands for student’s t-test between the specific nucleoporin and pooled data of all core nups together, no *p*-value is indicated if the difference is not significant; the sample size per strain is indicated (n).

## Results and discussion

### Circle-bound NPCs lack the nuclear basket

In order to study whether the anchorage of DNA circles to NPCs affects the NPCs’ composition, we took advantage of an engineered reporter DNA circle, which we have previously characterized (Baldi et al., 2017; Denoth-Lippuner et al., 2014; Shcheprova et al., 2008). The non-centromeric reporter DNA circle can be generated in vivo in a controlled manner. This circle is formed upon the induced excision of the centromere from a circular mini-chromosome and carries an array of 256 TetO repeats (Megee and Koshland, 1999). Hence, it is visible as a fluorescent dot in cells expressing TetR fused to a fluorescent protein (Fig. 1B, see methods and (Denoth-Lippuner et al., 2014). This circle replicates in S-phase and remains confined in the mother cell upon cell division, where it accumulates as the cell ages through successive rounds of cell division. Within 8-10 division cycles, the accumulating DNA circles cluster together to form a bright patch at the nuclear periphery. The attached NPCs accumulate with them in the nuclear envelope adjacent to the cluster (Denoth-Lippuner et al., 2014). Upon fusion of GFP to core nucleoporins, these NPCs are visible as a cap of enhanced fluorescence density (Fig. 1C, Figure 1-supplement 1A), different from young cells without accumulation of DNA circles, where nucleoporins are rather homogenously distributed in the nuclear envelope (Figure 1-supplement 1B). In the few cells that accumulate the circle in the population, this systems conveniently allows imaging circle-bound NPCs at single cell level, to discriminate them from unbound NPCs in the remainder of the same nuclear envelope.

Thus, we used this engineered DNA circle to quantify the local enrichment of single nucleoporins in the NPC cap compared to the rest of the nuclear envelope, to study the composition of DNA circle bound NPCs. We tagged 17 nucleoporins with GFP, representative of different NPC subcomplexes, in cells where the DNA circle cluster was labelled in red, owing to the expression of TetR-mCherry (Fig. 1B, C). In these images, all the stable core Nups, i.e. the scaffold components (outer ring: Nup84, Nup133; inter ring: Nup170, Nic96) and the components of the transport channel (FG-Nups: Nup53, Nup59) accumulated in the cap to similar extents. Quantification of their indicated a median of 2.4-fold enrichment, compared to their localization elsewhere in the nuclear envelope; Fig. 1D, E). All tested core nucleoporins showed the same enrichment in the cap covering the DNA circles, indicating that the circles bind intact NPC cores. The enrichment level of core Nups in the cap served therefore as reference to determine whether peripheral Nups were stoichiometrically associated with core NPCs bound to circles or depleted from those pores.

In striking contrast to core Nups, most peripheral subunits on both sides of the NPC were found to not accumulate in the cap. Four out of the five components of the nuclear basket and four out of the six components of the cytoplasmic complexes were excluded from the cap (Fig. 1C, E). Only Nup1, a stable nucleoporin at the nuclear side, and Nup159 and Nup83, which form the docking site for the cytoplasmic complexes at the cytoplasmic side of NPCs, accumulated together with core Nups. The basket components Mlp1 and 2, Nup60 and Nup2, and the subunits at the cytoplasmic side Nup116, Nup42, Gle1 and Gle2 were extensively excluded from the cap. The exclusion was the strongest for the basket proteins Mlp1 and Mlp2 (Fig. 1E), similarly to what is observed in the vicinity of the nucleolus (Galy et al., 2004), although the cluster of circles and the nucleolus do not overlap (Denoth-Lippuner et al., 2014). Altogether, these quantifications indicate that relative to the unbound NPCs, the circle-bound NPCs were specifically stripped of their nuclear basket and cytoplasmic complexes.

This picture is highly reminiscent of recent proteomics data indicating that Nup116, Nup2 and Nup60 are extensively lost from old yeast cells (Rempel et al., 2019). This study also identified Nsp1, a component of both the core and the cytoplasmic complex of the NPC (Bailer et al., 2001), as being lost with age. Because tagged Nsp1 is partially defective (Rajoo et al., 2018), we did not characterize its distribution. Thus, our results support previous observations indicating that old cells lose specific parts of the NPC, and suggest that this loss is driven by the attachment of DNA circles to the affected NPCs. Since Mlp1 and Mlp2 are required for DNA attachment to NPCs (Texari et al., 2013), including the attachment of DNA circles (Shcheprova et al., 2008), these proteins might mediate the recruitment of the DNA to NPCs but be subsequently displaced from them. Furthermore, our observations suggest that the loss of NPC components is not limited to those picked up by mass spectrometry but can be extended to nearly the entire basket and cytoplasmic complexes, including Mlp1 and Mlp2, Nup42, Gle1 and Gle2. Indeed, proteomic studies do not have the subcellular resolution obtained with our approach.

### The basket and cytoplasmic complexes are displaced from NPCs in wild type cells aging under physiological conditions

To evaluate whether remodeling of the circle-bound NPCs was indeed reflecting what happens in cells undergoing unperturbed aging, we next asked whether the basket and cytoplasmic complexes dissociates from NPCs of old cells not carrying our reporter circles. For some reason, accumulating ERCs are rather dispersed throughout the nuclear periphery and form clusters only episodically, such that NPC caps are less prominently observed in these cells. Clustering of the reporter circle might be stabilized by the fluorescent label (mCherry still has a low affinity for itself) or ERCs have means to escape clustering. In order to study if NPC remodeling happens under these conditions, we co-labelled pairs of Nups with distinct fluorophores and characterized their co-incorporation into NPCs by analyzing the spatial correlation of their fluorescence in the nuclear periphery as the cells aged (Fig. 2A). Indeed, the signals of the two labelled Nups should correlate well with each other as long as both colocalize to NPCs, and poorly if any one of them is displaced from NPCs (Fig. 2A).

**Figure 2.**
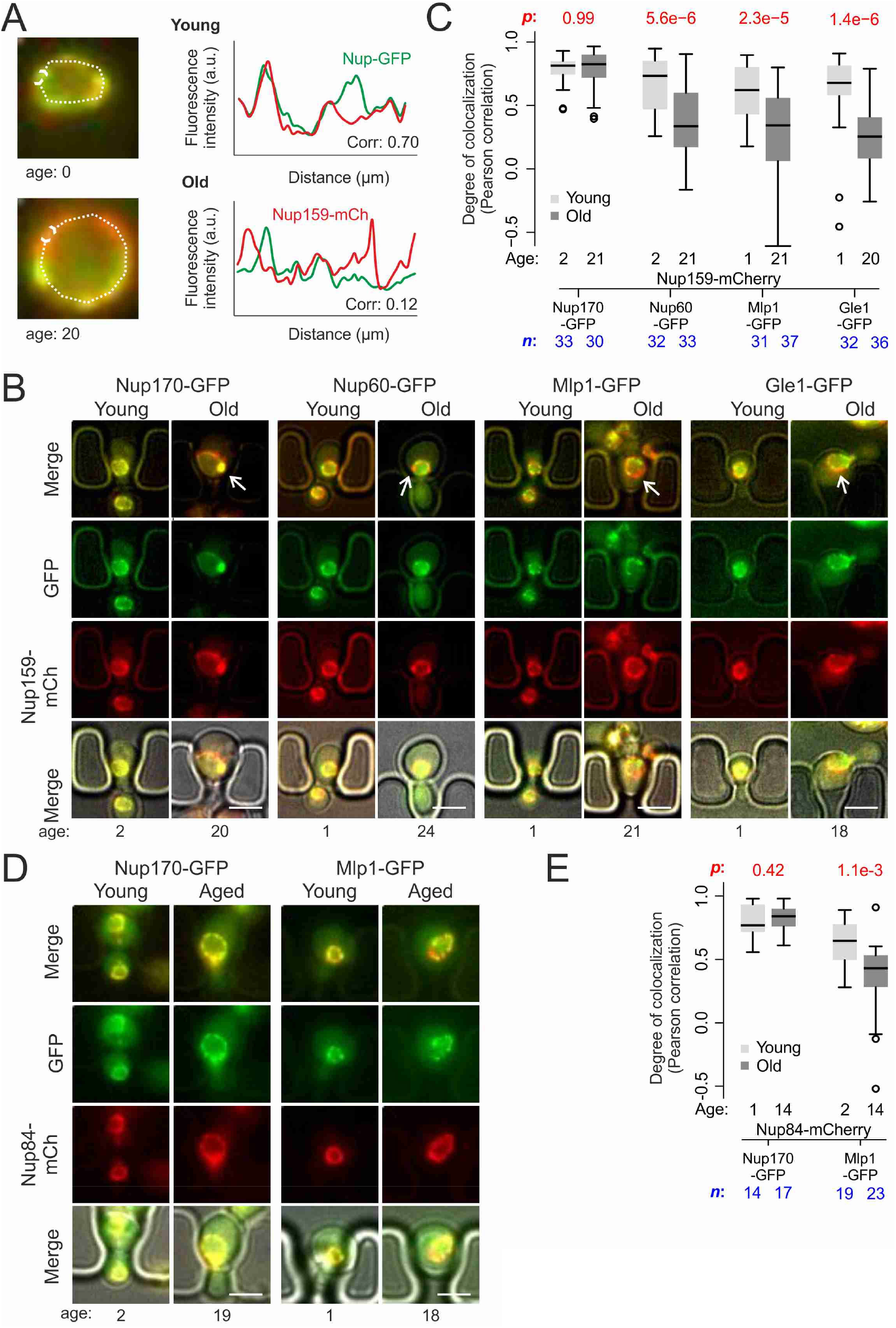
Peripheral subunits of NPC displaced in wild type aged cells. (A) To quantify the degree of nucleoporin colocalization, a line was drawn through the nuclear envelope in a focal image of a young (top) and aged (bottom) nucleus of cells co-expressing GFP-tagged Nup GFP (e.g. Nup60-GFP) and a mCherry-tagged Nup as a reference (e.g. Nup159-mCherry, see material and methods). The Pearson correlation between the two intensity profiles is calculated, and used as a measure for nucleoporin colocalization. (B) Fluorescent images of young and old cells in the yeast aging chip, with nucleoporins labeled with GFP (green) and the reference Nup159 with mCherry (red). The age of each cell is indicated. Scale bars are 5 µm. (C) Quantification of the degree of colocalization between target and reference nucleoporin is plotted for young and old wt. The *p*-value stands for the student’s t-test between young and old cells. The sample size (*n*) and the median age is indicated. (D) Fluorescent images of young and aged cells in the yeast aging chip, as in A, but with Nup84- mCherry as a reference nucleoporin. Scale bars are 5 µm. (E) Quantification of the co-localization between nucleoporins in young and old cells, as in D, but with Nup84-mCherry as a reference.

In order to acquire images of old mother cells, the labelled cells were loaded on a microfluidic chip (Jo et al., 2015) and imaged for 26 hours (21 divisions in average). A continuous flow of fresh medium provided nutrients to the trapped mother cells and flushed their daughter cells out, allowing the continuous imaging of the isolated mother cells. Bright field images were taken every 15 minutes to monitor budding events and record the replicative age of each cell. Fluorescence images were acquired after 2 and 26 hours (i.e. on average 2 and 21 divisions, respectively, Fig. 2B) and the degree of colocalization between two Nups was quantified at these two time points (Fig. 2C).

Analysis of the Pearson correlation between the signals of two core nucleoporins, namely Nup159 (labelled with mCherry) and Nup170 (labelled with GFP), showed that it was high in young cells and remained similarly high in old yeast mother cells (average correlation = 0.79±0.02, *n*=30 and 0.78±0.03, *n*=33, respectively; Fig. 2C). Thus, these two proteins remain indeed stable at NPCs of old cells. In contrast, the basket proteins Nup60 and Mlp1 and the cytoplasmic protein Gle1 (labelled with GFP) colocalized less extensively with Nup159 already in young cells (average correlation = 0.68±.0.04, *n*=32; 0.61±0.04, *n*=31 and 0.62±0.05; *n*=32 respectively), consistent with these proteins being more mobile and transiently associated with NPCs (Denning et al., 2001; Denoth-Lippuner et al., 2014; Dilworth et al., 2001; Niepel et al., 2013). More importantly, the correlation substantially dropped further in old cells (average correlation 0.35±0.05, *n*=33; 0.31±0.06, *n*=37 and 0.24±0.06, *n*=35 respectively; *p*<10^-4^; Fig. 2C). A similar drop of correlation was observed when comparing the signals of Mlp1-GFP and Nup82-mCherry (Fig. 2D, E), indicating that this correlation drop did not depend on the core Nup used as reference. Thus, the basket and cytoplasmic complexes are indeed displaced from a substantial fraction of the NPCs in old cells, upon unperturbed aging.

### The ERCs mediate the displacement of the basket from NPCs of old mother cells

To determine whether this remodeling of the NPC upon aging is driven by the accumulation and attachment of endogenous DNA circles, we first asked whether delaying ERC accumulation also delayed the removal of the basket from NPCs. Thus, we deleted the *FOB1* gene, which promotes ERC formation, and tested whether this restored the presence of the basket components Nup60 and Mlp1 at NPCs of old mutant cells (Fig. 3A). Strikingly, no significant dissociation of the basket from NPCs was observed in old *fob1Δ* mutant cells compared to wild type at the same age (same number of divisions) or to young *fob1Δ* mutant cells (Fig. 3A). Moreover, deleting the *SGF73* gene, which docks SAGA to NPCs and mediates circle anchorage to nuclear pores, also restored the localization of the basket to NPCs in aged cells (Fig. 3B). Thus, ERC presence and attachment is required for the basket to be displaced from NPCs.

**Figure 3.**
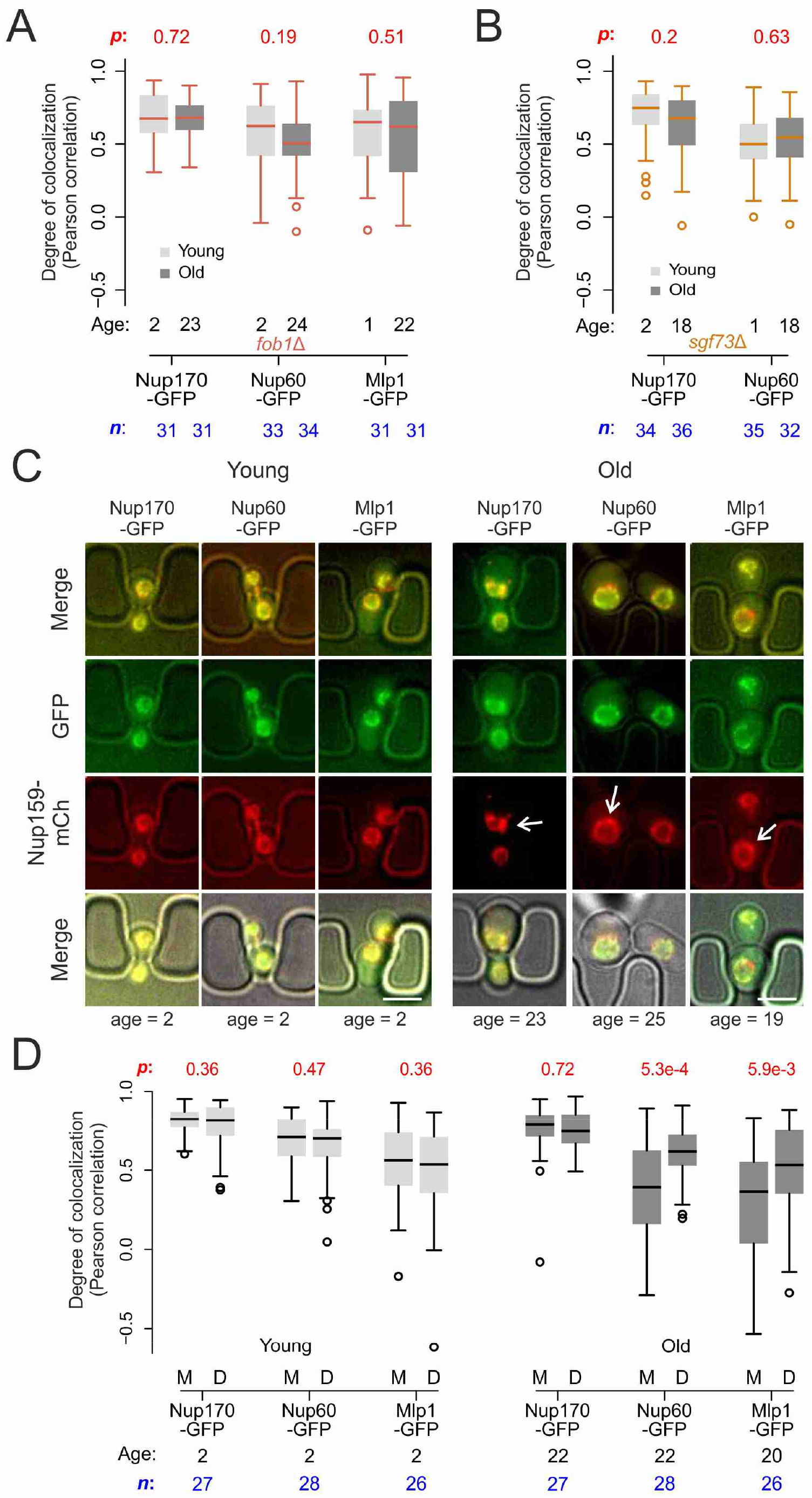
Endogenouse DNA circles drive nuclear basket displacement in wild type aged cells. (A) Quantification of the degree of colocalization between target and reference nucleoporin is plotted for young and old wt, as was done in Fig. 2C, but in *fob1Δ* cells. The sample size (*n*) and the median age is indicated. The *p-*value stands for the student’s t-test between young and old cells. (B) Same as A, but in *sgf73Δ* cells. (C) Fluorescent images of young and old mitotic cells in the yeast aging chip. Scale bars are 5 µm. (D) Same as in Fig. 2C, but comparing nucleoporin colocalization in young and old mother (M) with their corresponding daughter (D) cell. The *p-*value stands for the student’s t-test between mother and daughter cell. The sample size (*n*) and the median age is indicated.

Finally, as ERC anchorage displaces the basket from a substantial fraction of the NPCs in old wild type mother cells, their daughters, which do not inherit ERCs are expected to rapidly restore the proper localization of the basket proteins. Indeed, when comparing signal correlation between the basket proteins Nup60 or Mlp1, with the core Nup, Nup159, colocalization between basket and core nucleoporins was extensive in the rejuvenated daughter cells of old mothers, similar to what is observed in both the young mothers and their daughters (Fig. 3C, D). The correlation between basket and core Nups was only reduced in aged mothers. Altogether, we conclude that the formation of ERCs and their attachment to NPCs drives the displacement of the nuclear basket from pores. ERC accumulation is a direct cause for the displacement and loss of the nuclear basket from NPCs in physiologically aging yeast cell. This event probably mediates the subsequent displacement and loss of the cytoplasmic complexes (see below).

### The cells do not recognize remodeled NPCs as defective

Several studies have indicated that NPCs can deteriorate with time and established the existence of a machinery dedicated to removing damaged or misassembled NPCs. This quality control system recruits the ESCRT III machinery, including the adaptor protein Chm7, to defective NPCs and mediates their removal from the nuclear envelope (Rempel et al., 2019; Webster et al., 2016). Thus, we next asked whether the attachment of DNA circles and basket removal induces a defect that would be recognized as damage by the cell. To address this question, we investigated whether the circle-bound NPCs recruit Chm7 more frequently than bulk NPCs. We recorded the localization of Chm7, tagged with GFP, in cells loaded with DNA circles and asked whether it accumulated in the cap (Fig. 4A). The cells containing a cluster of DNA circles were categorized in the following classes: 1) the cells showing at least one Chm7 focus in the NPC-cap, 2) those showing a Chm7 focus somewhere else in the nuclear envelop and 3) those showing no visible Chm7 foci in the nuclear envelope (Fig. 4A). Interestingly, most cells formed either no Chm7 focus at the nuclear periphery (category 3, 41% of cells, *n* = 190 cells with circle cluster) or the focus formed was not associated with the cluster of DNA circles (category 2, 23% of the cells). Only about a third of the cells (category 1, 36%) formed a Chm7 focus adjacent to the circle-cluster, i.e. where the NPC-cap is located. Note that these Chm7-labeled foci are much smaller than the NPC-cap (Fig. 1C, 4A), indicating that if any, only few of the NPCs in the cap are targeted by this machinery. Importantly, based on fluorescence intensity measurements using Nup84-GFP in DNA circle loaded cells (Fig. 1C), we estimate that about 45% (+/- 12%, *n* = 10) of the NPCs is sequestered in the NPC-cap at that stage. Thus, this analysis did not reveal any substantial enrichment of Chm7 overlapping with the clusters of DNA circle; the occurrence of Chm7 in the NPC cap seemed to be rather coincidental. We concluded that the circle- bound NPCs are not detected as defective by the cell more or less than the other NPCs in the rest of the nuclear envelope.

**Figure 4.**
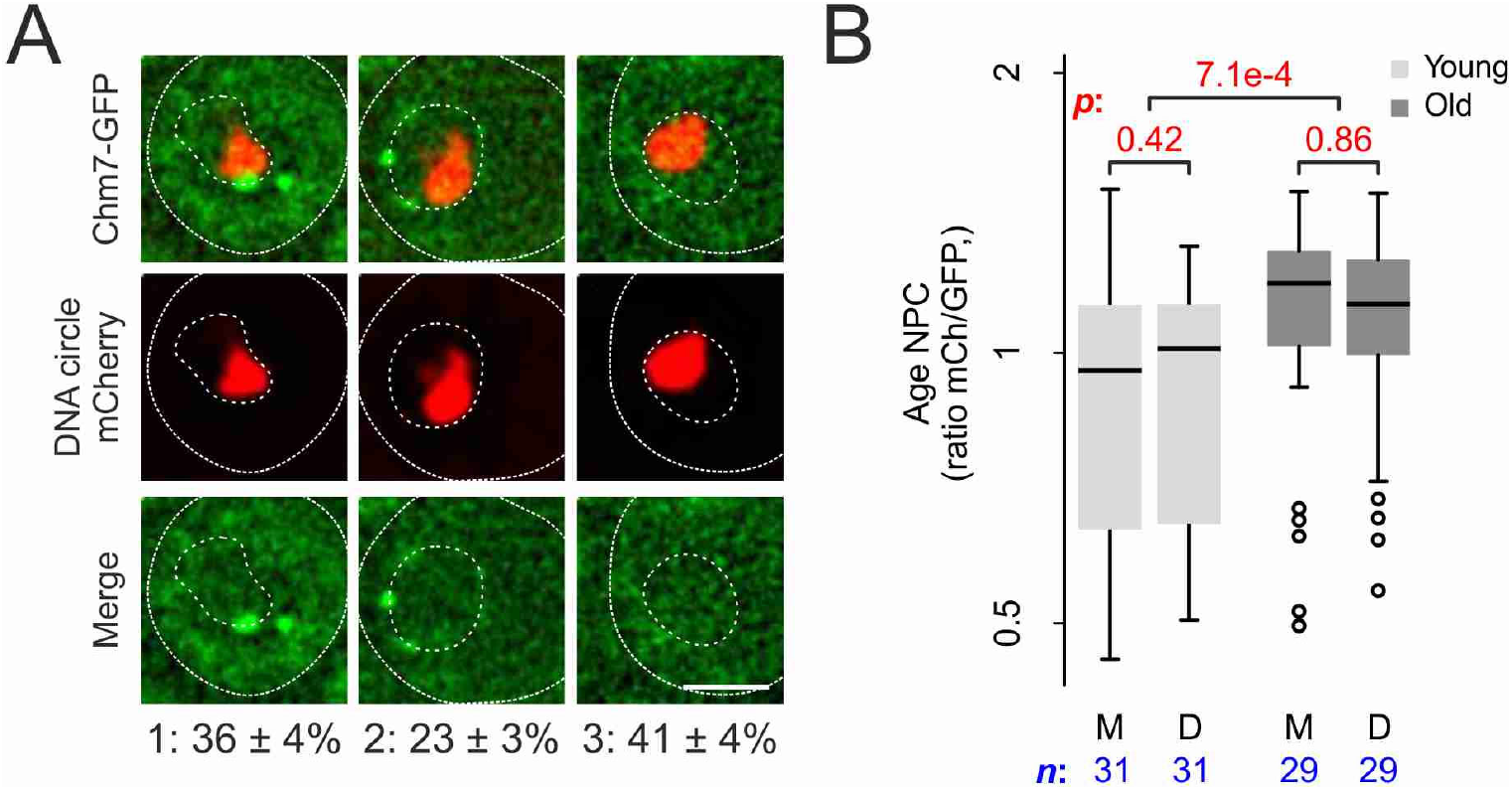
DNA circle anchored NPCs are not recognized as defective. (A) Fluorescent images of nuclei in yeast cell with accumulated DNA circles labeled with TetR-mCherry and Chm7-GFP. Occurance of Chm7-GFP localization is categorized 1) at least one Chm7 focus in the NPC-cap, 2) a Chm7 focus somewhere else in the nuclear envelop and 3) no visible Chm7 foci. The standard error of the mean is indicated, *n* = 190. (B) The relative age of the NPC plotted as the intenstity ratio between mCherry and GFP of Nup170 tagged with a fluoresecent timer (Nup170-mCherry-sfGFP), measured in young and old mother (M) versus daughter cells (D).

### The remodeled NPCs are not particularly old

Since the anchorage of a DNA circle to an NPC causes its retention in the mother cell, we next wondered whether this could cause the progressive accumulation of older NPCs in aged mother cells. To test this possibility, we measured the relative age of NPCs in old mothers and their daughter cells using the tandem fluorescent protein timer, consisting of mCherry (mCh) and superfolder GFP (sfGFP) (Khmelinskii et al., 2012). Due to different maturation kinetics between the two fluorophores, a newly synthesized protein appears first in the green channel before acquiring the red fluorescence over time. As the turnover rate of Nup170 is very low in NPCs (D’Angelo et al., 2009), older pores with tagged Nup170 are expected to emit more red fluorescence than green in comparison to newly assembled pores. To see if old mothers are enriched in red-shifted old pores, we loaded the cells expressing Nup170-mCh-sfGFP on the microfluidic chip and imaged them as above (Fig. 2B) as the cells aged (Jo et al., 2015). The fluorescence channels were recorded after 2 and 26h. We did observe a tendency for young cells to put slightly more red shifted NPCs in the bud than in the mother cell (Fig. 4B, not statistically significant, *n*=31 mother-bud pairs), as was shown before (Khmelinskii et al., 2012). Over time we observed a highly significant increase of the red fluorescence signal relative to the green signal in old compared to young cells (*n*=29 old cells, p<0.001), indicating that old cells actually accumulate old pores over time. Although we observed a trend for the NPCs of the daughters of old mother cells to be slightly younger, indicative for some retention of old pores in the mother cell, this difference was not significant in our analysis (Fig. 4B).

Thus, together these data make three points. First, the data support earlier findings indicating that pre-existing NPCs in young cells are not particularly retained in the mother cell and even that their segregation might be biased towards the daughter cell (Khmelinskii et al., 2012). Second, the data establish that over time the mother cell does accumulate older NPCs, possibly with the accumulation of ERCs. Third, these older NPCs are however not tightly retained in the old mother cell. As circle- bound NPCs are retained in the mother cell (Fig. 3C, D, (Denoth-Lippuner et al., 2014)), these findings suggest that old NPCs do exchange the circles with younger ones. This conclusion is in agreement with the results from photobleaching experiments previously reported (Denoth-Lippuner et al., 2014), indicating that although NPC exchange is slow between the cap and the rest of the nuclear envelope, it does happen. Thus, we concluded that although circle-bound NPCs accumulating in mother cells are rather old in average, they are not significantly older than those of the rejuvenated daughter cells. Thus, together our data do not support the notion that the age of the old mother NPCs drives their remodeling.

### Nucleoporin acetylation promotes NPC remodeling

Since DNA circles attach to NPCs via the SAGA complex, we next wondered whether basket removal could be driven by their SAGA-dependent acetylation. Indeed, three of the four basket proteins displaced from NPCs upon circle attachment are acetylated *in vivo*, and the acetylation of Nup60 and Nup2 is mediated at least in part by SAGA (Downey et al., 2015; Henriksen et al., 2012). Testing whether removing SAGA activity altogether restored the recruitment of the basket proteins was not possible in our “cap” assay since knocking out its catalytic subunit, encoded by *GCN5*, abolished circle clustering and cap formation (Fig. 5A), consistent with SAGA and its acetyl-transferase activity mediating circle anchorage to NPCs (Denoth-Lippuner et al., 2014).

**Figure 5.**
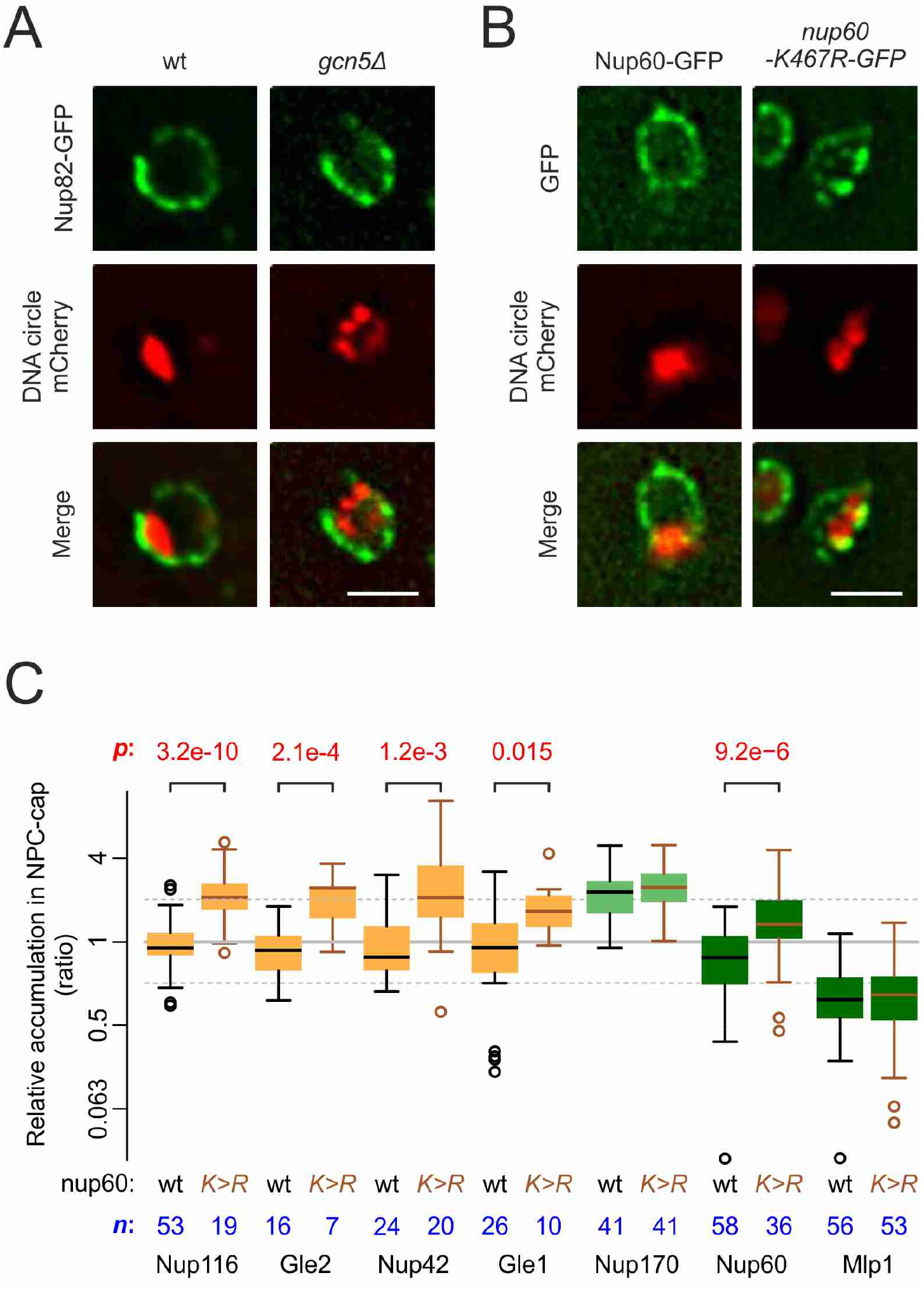
Basket displacement is post-transcriptional regulated. (A) Example images of nuclei in yeast cells with accumulated DNA circles and Nup82-GFP in wt and *gcn5Δ* cells. The DNA circle clusters are labeled with TetR-mCherry (red). Scale bar is 2 µm. Same as A, but with Nup60-GFP and nup60-K467R-GFP. (C) Quantification of GFP-labeled nucleoporin accumulation in the cap, in nup60- K467R (brown lines), compared to wt (black lines) on a log2-scale, as in Fig. 2B. Wt data is a copy from Fig. 2B. The *p-*value stands for the student’s t-test between accumulation ratio of a specific nucleoporin in wt and *nup60-K467R*, no *p*-value is indicated if the difference is not significant. The sample size (*n*) is indicated.

Thus, instead of inactivating SAGA we asked whether acetylation of the basket proteins contributes to displacing them from pores. Nup60 is prominently acetylated on lysine 467(Choudhary et al., 2014, 2009). Therefore, we asked as a proof of principle whether preventing this modification by substituting lysine-467 by arginine was sufficient to abrogate the exclusion of Nup60 from the cap (Fig. 1C, E). Strikingly, not only Nup60-K467R now accumulated in the cap nearly as much as core nucleoporins, but the cells expressing this protein also restored the localization of the cytoplasmic complexes (Nup116, Nup42, Gle1 and Gle2) to cap NPCs (Fig. 5B, C). In contrast, the basket component Mlp1 remained displaced, indicating that the anchorage of DNA circles might also lead to other modifications in different basket proteins, independently of that of Nup60 at lysine-467. Thus, we concluded that the acetylation of Nup60 on K467 upon circle attachment drives Nup60’s displacement from NPCs and subsequently that of the cytoplasmic complexes as well. How Nup60’s presence at the nuclear pore regulates the recruitment or stabilization of the cytoplasmic complexes on NPCs is not known at this stage. Possibly a signal is transduced from the nuclear to the cytoplasmic side of the NPC, or the clustering of the pores could drive the displacement of the cytoplasmic complexes. Furthermore, additional events displace the Mlp1/2 proteins as well, possibly also through their own acetylation or that of Nup2.

Thus, altogether we found no indication for the anchorage of DNA circles causing NPC deterioration. Rather, our data indicate that circle anchorage promotes the displacement of the nuclear basket through post-translational modifications, and that this in turn causes the detachment of the cytoplasmic complexes. Considering the fact that Nup60 acetylation contributes to the regulation of gene expression (Kumar et al., 2018), and that the residence of the basket to NPCs is dynamic also in young cells, the removal of the basket upon circle attachment might reflect a common physiological process taking place each time chromatin interact in a SAGA-dependent manner with interphase NPCs. The fact that DNA circles remain attached to NPCs during mitosis, whereas chromosomes detach from the nuclear periphery at that stage (Kanoh, 2013), might be caused by their inability to undergo condensation and recruit the deacetylase Hst2 (Kruitwagen et al., 2018). Further studies will be needed to determine the role of acetylating other Nups beyond Nup60 in basket displacement, circle anchorage and aging, and whether Hst2 reverts the acetylation of these proteins at chromosome-attached NPCs during mitosis.

### Basket displacement promotes aging

Thus, together our results indicate that DNA circles modulate the organization of NPCs as they attach to them, leading to the accumulation of remodeled NPCs as circles accumulate and the cell progresses in replicative aging. Therefore, we wondered whether the effect of DNA circles on NPC organization is at least a part of the mechanisms by which ERCs promote cellular aging. Thus, we next investigated whether interfering with NPC remodeling has an impact on the lifespan of the cells. We took advantage of a new microfluidic chip design, that could retain more efficiently cells during their entire lifespan (i.e. >95% of the cells were kept until cell death (Fig. 6A, B), see methods). When monitoring wild type cells in this setup, their replicative life span was reproducibly of 18 generations (Fig. 6C). Although this lifespan is relatively low, it is within the range of RLSs reported for wild type cells measured on different microfluidic platforms (Chen et al., 2017; Janssens and Veenhoff, 2016). Interestingly, mutant cells with pores lacking the central basket component Nup60 aged rapidly (median life span of 13 divisions), while the *nup60-K467R* mutant cells already showed an extended longevity (20 divisions, Fig. 6C), despite the proteins Mlp1 and Mlp2 remaining displaced in these cells. Thus, our observations support the idea that displacement of the basket by SAGA promotes aging, while basket stabilization fosters longevity. Together our data indicate that the remodeling of NPCs upon DNA circle attachment to them is one causal element for how ERC accumulation limits the longevity of yeast mother cells.

**Figure 6.**
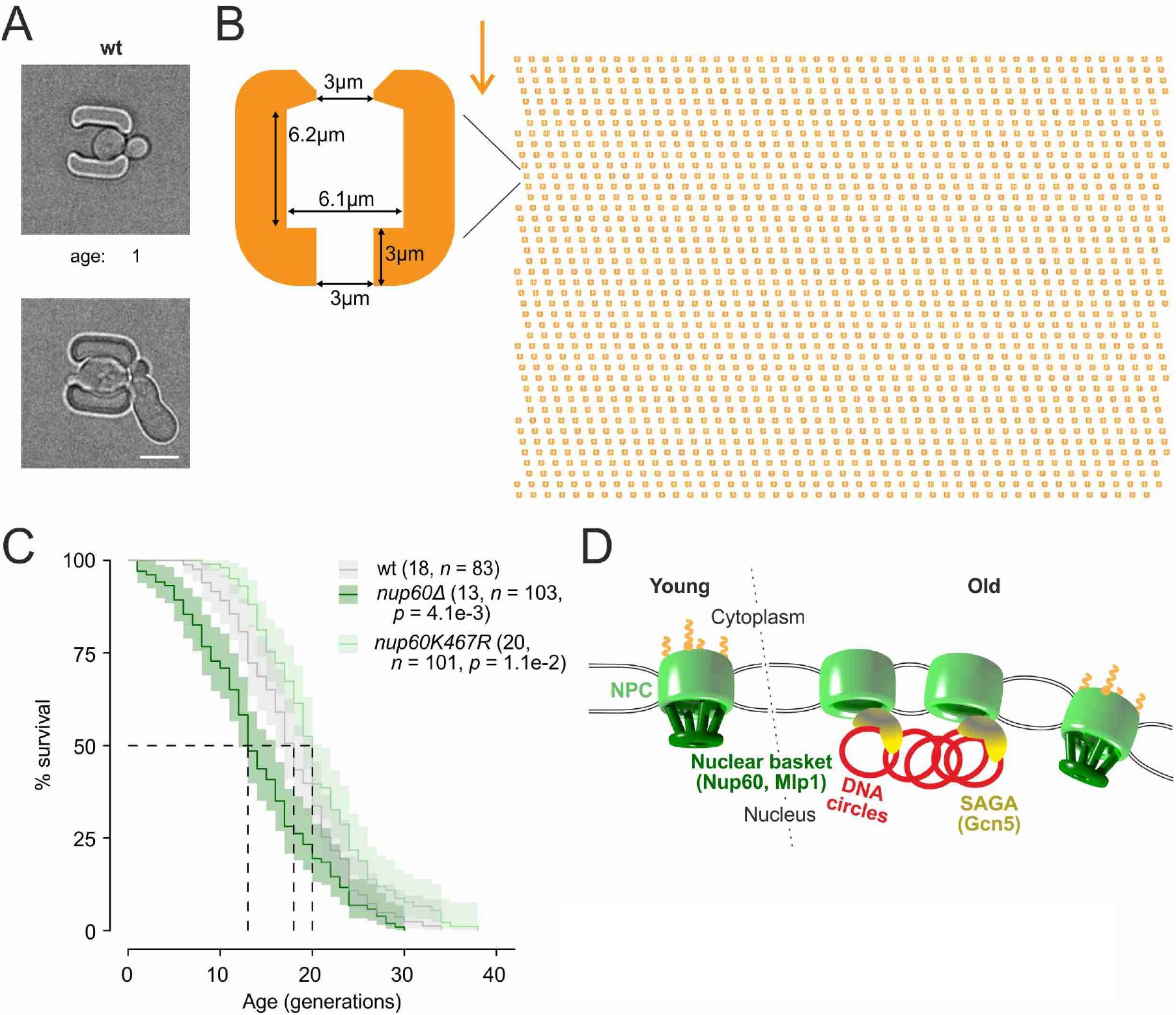
Basket displacement promotes aging. (A) Example images of the same young and old wt cell in a trap of the yeast aging chip. The age of the cell is indicated, scale bar is 5 µm. (B) The design of the array with improved traps in the aging chip. One single trap is highlighted to show its dimensions. The arrow shows the direction of the medium flow. (C) The lifespan curve for *nup60Δ* and *nup60-K467R* compared to a wt strain, plotted with 95% confidence interval limits. The *p-*value stands for Log-Rank test between *nup60Δ* or *nup60-K467R* with the wt strain. (D) Model depicting that DNA circle accumulation affects the organization of the NPC via a SAGA-mediated basket displacement.

Strikingly, the displacement of the basket from NPCs is highly reminiscent of what has been already described for the NPCs adjacent to the nucleolus in yeast. Indeed, the yeast nucleolus adheres to the nuclear envelope and the NPCs bordering the nucleolus lack Mlp1/2 (Galy et al., 2004). The functional relevance of this observation is unknown but has suggested that these NPCs might be functionally specialized. Thus, we suggest that the accumulation of basket-less NPCs with age does not reflect an accumulation of defective NPCs per-se, but rather leads to an imbalance of specialized over non-specialized NPCs in old cells (Fig. 6D).

### mRNA export and mRNA surveillance factors are specifically displaced from circle-bound NPCs

Thus, we reasoned that NPC specialization might have some regulatory effect on their function. Nuclear pore complexes mediate the transport of cargos between nucleoplasm and cytoplasm and therefore transport factors transiently localize to NPCs in young and healthy cells (Derrer et al., 2019; Kumar et al., 2002). Any effect of NPC remodeling on the localization of these factors to NPCs may reflect changes in their dynamics within NPCs and hence, on NPC functionality. Therefore, we characterized how circle anchorage affected the recruitment of transport factors and other associated proteins to NPCs. We labeled a broad panel of transport and associated factors with GFP and quantified their nuclear localization in respect to the DNA circle cluster, as in Fig. 1. Only the factors showing a clear (transient) localization to the nuclear envelope of young wild type cells were characterized. Consistent with circle-bound NPCs lacking a basket, the two basket-associated proteins Esc1 and Ulp1 (both involved in telomeric silencing and mRNA surveillance, (Bonnet et al., 2015)) were excluded from NPC caps (Fig. 7A, B). In striking contrast, none of the 10 importins tested were displaced from the cap (Fig. 7B). Likewise, the exportins Xpo1 (Crm1), Msn5 and Los1, which ensure the export of proteins and tRNAs, accumulated with the cap to similar extent as the core Nups. Since nearly all these proteins accumulated to the same extent as core NPCs in the cap compared to elsewhere in the nuclear envelope, we concluded that basket-less pores interact with them with similar dynamics as bulk NPCs. Two importins were in average significantly further enriched at the cap, namely Srp1/Kap60 and Kap123. This may indicate either that these two nucleoporins shuttle more intensely through basket-less NPCs, or that they linger longer in them. Strikingly, Kap123 mediates the nuclear import of ribosomal proteins, promoting their subsequent assembly into ribosomes, and the import of histones H3 and H4. Both ribosome and nucleosome assembly have been shown to become reduced in aged yeast cells (reviewed in (Matos-Perdomo and Machín, 2019)).

**Figure 7.**
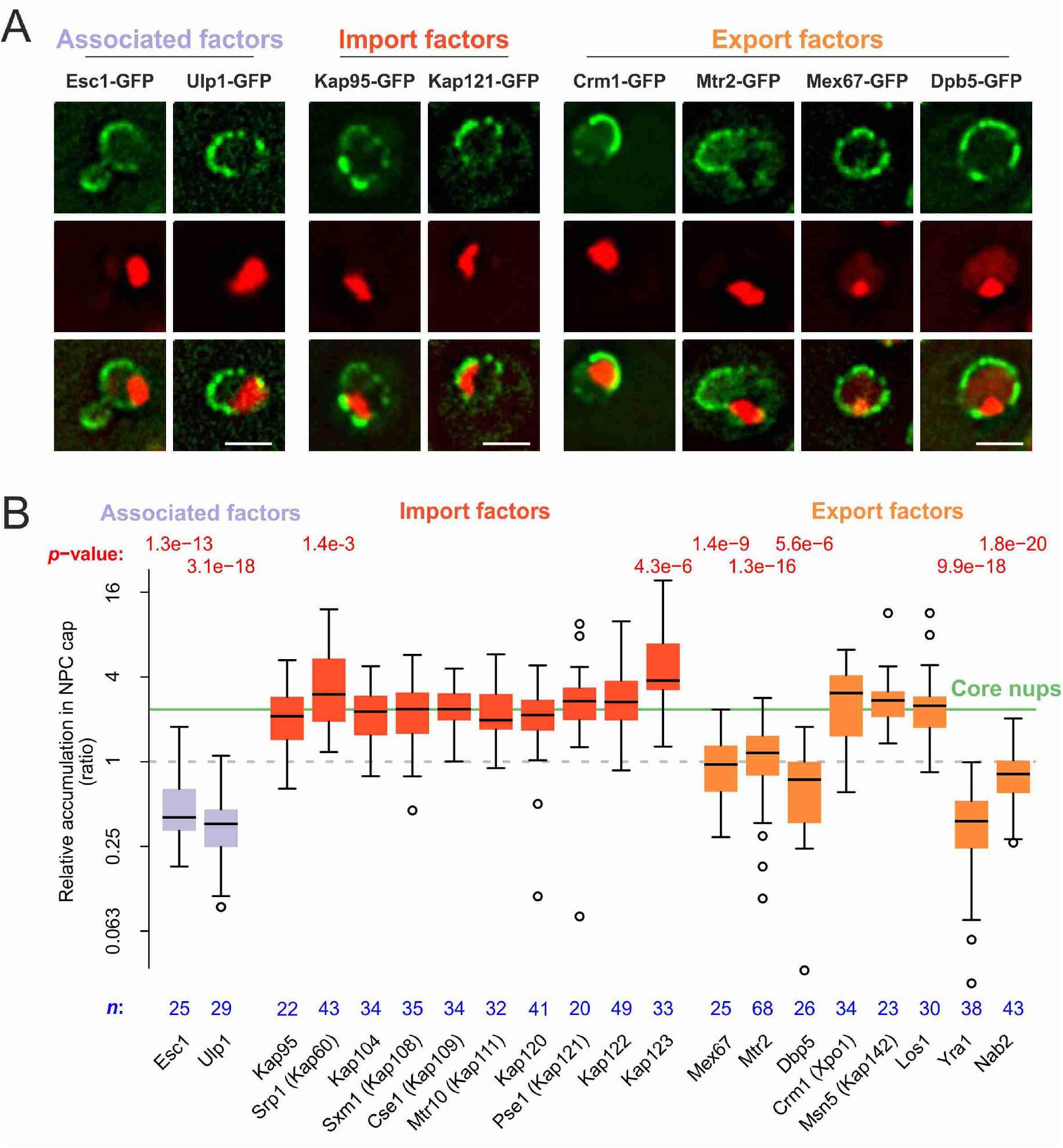
mRNA export and surveillance factors are displaced from circle-bound NPCs. (A) Fluorescent images of nuclei in aged yeast cells, where the DNA circle clusters are labeled with TetR-NLS-mCherry (red) and transport factors with GFP (green). Scale bar is 2 µm. (B) Quantification of factor recruitment in the NPC cap adjacent to the DNA circle cluster, as in Fig. 2B. The protein accumulation is plotted for associated factors (grey), import factors (red) and export factors (orange), and is compared to accumulation of the pooled NPC core subunits NPC (green line, duplicated from Fig. 2B), no *p*-value is indicated if the difference is not significant. The sample size (*n*) is indicated.

More strikingly, 5 exportins were specifically depleted from the cap (Fig. 7B), indicating that their interaction with core NPCs was substantially decreased. Interestingly, all of these exportins are involved in mRNA export. Thus, all seven proteins that we find to be excluded from the cap are involved in this process (Bonnet et al., 2015; Iglesias et al., 2010; Stewart, 2010). We postulate that circle-bound NPCs are specifically inhibited for their function in mRNA export.

### Conclusion

In summary, our data indicate that old yeast cells accumulate an increasing proportion of NPCs depleted of the nuclear basket and cytoplasmic complexes and that may have a reduced capacity for mRNA export. Once the proportion of basket-associated NPCs goes below some threshold, the resulting imbalance become deleterious for the cell. Importantly, our study establishes that the stoichiometry changes observed with age are not due to NPC deterioration but rather to their specialization as they become increasingly decorated with DNA circles, mainly ERCs. The function of this specialization is not known but observed also in young cells, for example near the nucleolus.

Whether nucleolus-associated NPCs are depleted of other factors, such as the cytoplasmic complexes, has not been reported but our study suggests that it might be the case. Furthermore, the dissociation of the basket from NPCs appears to have broad functions beyond nucleolar NPCs in young cells since the acetylation of Nup60 is involved in the regulation of diverse loci on chromosomes, such as the activation of cell cycle genes (Kumar et al., 2018), and the SAGA complex is involved in the rapid activation of many genes in response to environmental changes such as heat shock, inositol starvation or changes in the available carbon source (Huisinga and Pugh, 2004; Kremer and Gross, 2009). In many of these instances, SAGA-dependent regulation involves the recruitment of the target locus to the nuclear periphery and their anchorage to NPCs. Thus, the modifications observed at circle-bound NPCs reflect physiological changes that are common in young cells and amplified through ERC accumulation in old cells. Future studies will require to determine how this imbalance affects cellular viability.

The notion that the age-dependent remodeling of NPC is not due to damages but to regulatory steps has consequences for our understanding of the evolution of aging. Indeed, a leading theory for the apparition of the aging process in evolution is that it is the result of traits and processes that are selected for their selective advantage early in life, despite of deleterious effects later-on (Kirkwood and Rose, 1991). Antagonistic pleiotropy has been difficult to document beyond the well accepted idea that sparing on quality control and damage-repair liberates resources for the generation of progeny early in life, at the expense of longevity (Ackermann et al., 2007; Austad and Hoffman, 2018; Williams, 1957). The case of NPCs and their role in SAGA-dependent gene regulation might depart from this notion by suggesting that the trade-off here is not a matter of quality control but of adaptability. Indeed our data suggest that the remodeling of NPCs with age is a secondary consequence of the role of chromatin-NPC interaction in the rapid and strongly response of cells to environmental challenges. The mechanistic relevance of these interactions during gene regulation, sometimes referred as gene gating, as well as their effects on NPC activity and possibly other processes at NPCs, such as DNA repair (Strambio-De-Castillia et al., 2010), are not yet fully understood but their characterization is likely to shed light into why and how aging has emerged, at least in yeast. Furthermore, our study indicate that aged cells may offer a model of choice for studying how NPC specialization affects their function.

There is currently no evidence for DNA circles contributing to aging in other organisms but there is strong evidence that chromatin interaction with the nuclear periphery is affected by age in many cell types. Most remarkably, progerin, a progeriatric isoform of Lamin A and causing the Hutchinson- Gilford progeria syndrome in humans, affects the recruitment of heterochromatin to the nuclear periphery, and displaces Nup153 and TPR (the mammalian homologs of Nup60 and Mlp1/2) from the nuclear periphery (Balmus et al., 2018; Cobb et al., 2016; Kelley et al., 2011; Larrieu et al., 2018). Thus, we suspect that the effects of aging on NPCs and the role of NPCs in aging might be strikingly similar between yeast and mammals.

## Materials and methods

### Strains and plasmid

All the yeast strains and plasmids used in this study are listed in table S1 and are isogenic to S288C. GFP-tag and knock-out strains were generated using classical genetic approaches (Janke et al., 2004)). All cultures were grown using standard conditions, in synthetic drop-out medium (SD-medium; ForMedium, Norfolk, UK) or indicated otherwise, at 30°C.

The non-chromosomal DNA circle was obtained from the Megee lab (Megee and Koshland, 1999) and contains an array of 256 TetO repeats, the centromere is flanked by target site for the R- recombinase; the β-estradiol-inducible expression recombinase (from the genome) drives the excision of the centromere and converts the minichromosome into a non-chromosomal DNA circle ((Baldi et al., 2017; Denoth-Lippuner et al., 2014; Shcheprova et al., 2008). TetR-NLS-mCherry is genetically expressed and labels the TetO repeats; the NLS (nuclear localization signal) ensures its accumulation in the nucleus.

The nup60-K468R strain was generated with a one-step CRISPR-Cas9 method, based on (Laughery et al., 2015). The gRNA was designed with an online tool to identify an optimal guide RNA (gRNA) target site in Nup60 (http://wyrickbioinfo2.smb.wsu.edu/crispr.html). The donor DNA with the Nup60K467R mutation and the gRNA-encoded DNA were ordered as single stranded oligo’s (Microsynth AG, Balgach, Switzerland) and annealed. The gRNA-encoded DNA was recombined into the Cas9 expression vector pML104_Cas9_HygR in a one-step approach, to induce the Nup60K467R mutation simultaneously: SwaI-linearized vector, the gRNA-encoded double stranded oligo’s and the donor DNA were transformed all at once into a wild type yeast strain and plated on hygromycin selection medium. The expression vector with gRNA (pML104_Cas9_HygR_Nup60) was rescued from cell material, propagated in bacteria and confirmed by digest analysis and sequencing. Genomic DNA was extracted from single yeast clones and the presence of the point mutation was confirmed by sequencing. Positive clones were propagated on YPD to get rid of the expression vector.

### Microscopy

For fluorescent microscopy, yeast cells were precultured for minimally 24h in synthetic drop-out medium. 1 ml of cells from exponential growing cultures with OD<1 were concentrated by centrifugation at 1.000xG, resuspended in ∼5 ul of low fluorescent SD-medium, spotted on a round coverslip and immobilized with a SD/agar patch. The cells were imaged in z-stacks of 6 slices with 0.5 μm spacing, with a 100×/1.4 NA objective on a DeltaVision microscope (Applied Precision) equipped with a CCD HQ2 camera (Roper), 250W Xenon lamps, Softworx software (Applied Precision) and a temperature chamber set to 30°C.

To accumulated DNA circles in the nuclei of aging mother cells, yeast cells were pre-cultured for 24h in SD–URA at 30°C and then shifted to SD-LEU medium supplemented with 1 µM β-Estradiol (Sigma-Aldrich, St. Louis, MO), incubated for 16-18h at room temperature. The β-Estradiol induced expression of the recombinase and the excision of the centromere, see:(Denoth-Lippuner et al., 2014). To visualize DNA clusters, we use specifically 1×1 binning and made short time-lapse movies of 15 min, intervals for 5 min.

### Aging microfluidic platform

Nucleoporin colocalization and the tandem fluorescent protein timer analysis during aging were investigated using the high-throughput yeast aging analysis (HYAA) microfluidics dissection platform (Jo et al., 2015). The PDMS (polydimethylsiloxane) microchannel is made by soft-lithography and bonded on the 30 mm micro-well cover glass in the 55 mm glass bottom dish (Cellvis, CA, USA). For the lifespan analyses, a chip with a new cell trapping design was used (Fig. 6A, B), to ensure excellent retention of old cells (see below).

To start the experiment, yeast cells were pre-cultured for 24h in SD-full supplemented with 0.1% Albumin Bovine Serum (protease free BSA; Acros Organics, Geel, Belgium). Young cells from a exponentially growing culture were captured in the traps of the microfluidic chip; the chip was continuously flushed with fresh medium at a constant flow of 10 μl/min, using a Harvard PHD Ultra syringe pump (Harvard Apparatus, Holiston, MA, USA) with two or four 60mL BD syringes, with inner diameter 26.7 mm (Becton Dickinson, Franklin Lakes, NJ, USA). Bright field images were recorded every 15 min. throughout the duration of the entire experiment. To measure the nucleoporin colocalization or pre-mRNA translation, fluorescent images only after 2h, 12h, 26h or/and 50h. For imaging we used an epi-fluorescent microscope (TiE, Nikon Instruments, Tokyo, Japan) controlled by Micro-Manager 1.4.23 software (μManager, PMID 25606571), with a Plan Apo 60x 1.4 NA objective. For fluorescence illumination of the GFP and mCherry labeled proteins, a Lumencor Spectra-X LED Light Engine was used. Stacks of 7 slices with 0.3 μm spacing were recorded during. The age of the cell was defined by the number of daughter cells that emerged during the budding cycles.

For the nucleoporin correlation analysis, a cell of interest was manually selected if it stays in the focal plane in the bright field channel. Its age was determined and a segmented line was drawn through the nuclear envelope in an image in the focal plane, using Fiji/ImageJ 1.51n (Schindelin et al., 2012), and the intensity profiles were recorded for both fluorescence channels. The Pearson correlation between the intensity profiles was calculated and plotted in R (R Development Core Team, 2011).

For the tandem fluorescent protein timer analysis, late mitotic cells were selected after 2h and 26h incubation in the chip. Its age was determined and a segmented line was drawn through the nuclear envelope in an image in the focal plane, as described above. The average background corrected intensity for GFP and mCherry was calculated and plotted in R.

To obtain a reliable lifespan curves, the majority of the cells should be retained until cell death to prevent biasing the data. Although different microfluidic dissection platforms have been developed, it is still a challenge to reach high enough retention efficiency in the microfluidics chip for life span analysis. Here we used an improved design of yeast cell traps, having small “claws” at both sides, preventing the escape of bigger cells at higher age (Fig. 6A, B). This allowed us to retain >95% of the cells during their full lifetime. Only bright field images were recorded every 15 min. throughout the entire experiment of 70-80 hours. All cells in a field of view were analyzed, the replicative lifespan was determined for each single cell by counting the budding cycles before cell death.

## Acknowledgements

We thank Joachim Hehl, Tobias Schwartz and the light microscopy center of ETH Zürich (ScopeM) for microscopy support; Rodrigo Merayo Martínez, Irina Remple and Liesbeth Veenhoff for their help with running the microfluidic chips. We acknowledge financial support by Swiss National Science Foundation (grant 31003A-105904 to Y.B.).

## Figures

**Figure 1 -supplement 1.**
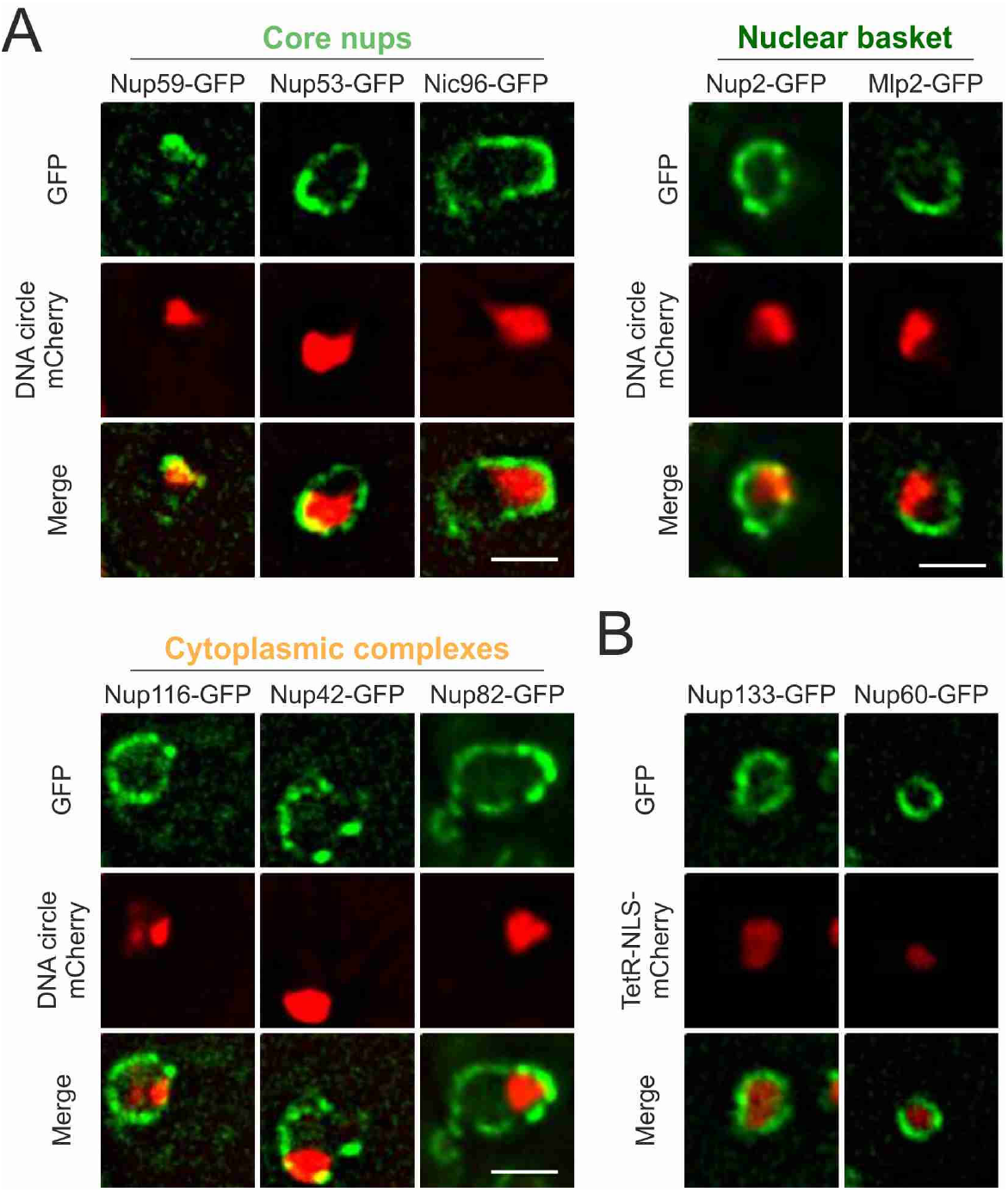
DNA circle anchored NPCs lack the peripheral subunits. , Related to figure 1. (A) Additional example images of all strains as quantified in Figure 1E. The DNA circle clusters are labeled with TetR-NLS-mCherry (red) and different nucleoporins with GFP (green). Scale bar is 2 µm.(B)Example images of nuclei in young yeast cells prior to DNA circle accumulation; with nucleoporins labeled with GFP and soluble TetR-NLS-mCherry. Without Tet-O containing DNA circles, TetR-NLS-mCherry is homogenously distributed in the nucleoplasm and the localization of both core subunits (Nup133-GFP) and basket subunits (Nup60-GFP) is rather homogenously distributed throughout the nuclear envelope.

